# Competition between mobile genetic elements drives optimization of a phage-encoded CRISPR-Cas system: Insights from a natural arms-race

**DOI:** 10.1101/381962

**Authors:** Amelia C. McKitterick, Kristen N. LeGault, Angus Angermeyer, Muniral Alam, Kimberley D. Seed

**Affiliations:** Department of Plant and Microbial Biology, University of California, Berkeley, 111 Koshland Hall, Berkeley, CA 94720, USA; International Centre for Diarrhoeal Disease Research, Bangladesh, Dhaka, Bangladesh; Chan Zuckerberg Biohub, San Francisco, CA 94158, USA

**Keywords:** CRISPR, phage, mobile genetic elements, cholera

## Abstract

CRISPR-Cas systems function as adaptive immune systems by acquiring nucleotide sequences called spacers that mediate sequence-specific defense against competitors. Uniquely, the phage ICP1 encodes a Type I-F CRISPR-Cas system that is deployed to target and overcome PLE, a mobile genetic element with anti-phage activity in *Vibrio cholerae*. Here, we exploit the arms race between ICP1 and PLE to examine spacer acquisition and interference under laboratory conditions to reconcile findings from wild populations. Natural ICP1 isolates encode multiple spacers directed against PLE, but we find that single spacers do not equally interfere with PLE mobilization. High-throughput sequencing to assay spacer acquisition reveals that ICP1 can also acquire spacers that target the *V. cholerae* chromosome. We find that targeting the *V. cholerae* chromosome proximal to PLE is sufficient to block PLE and propose a model in which indirect chromosomal spacers are able to circumvent PLE by Cas2-3-mediated processive degradation of the *V. cholerae* chromosome before PLE mobilization. Generally, laboratory acquired spacers are much more diverse than the subset of spacers maintained by ICP1 in nature, showing how evolutionary pressures can constrain CRISPR-Cas targeting in ways that are often not appreciated through *in vitro* analyses.

## Introduction

Phages often vastly outnumber their bacterial hosts in a variety of environments (1). As such, bacteria have evolved numerous mechanisms for phage defense, including adaptive immunity via clustered regularly interspaced short palindromic repeats (CRISPR) and CRISPR-associated (Cas) proteins (2,3). CRISPR-Cas systems are composed of a CRISPR array—a series of “spacers” of foreign sequence alternating with repeats that are transcribed into CRISPR RNAs (crRNAs)—and CRISPR-associated (Cas) genes. Together with crRNAs, Cas proteins defend against foreign nucleic acids, such as the genome of an infecting phage, through a three-step process: adaptation, crRNA expression, and interference. During adaptation, a foreign DNA fragment is incorporated into the CRISPR array to provide a molecular memory of the challenges that the host cell has faced. This CRISPR array is expressed and processed into individual crRNAs, which complex with Cas proteins and survey the cell for complementary invading nucleotides. Upon finding a complementary sequence, termed protospacer, a Cas nuclease is recruited to the site to mediate interference by cleaving the substrate, ultimately leading to the destruction of the invader (4,3). Across CRISPR-Cas containing bacteria and archaea, Class 1 Type I CRISPR-Cas systems employing a Cas3 enzyme for DNA unwinding and degradation (5), are the most prevalent (6).

CRISPR-Cas systems do not discriminate between horizontally acquired traits based on fitness gain or loss. Hence, CRISPR-Cas systems are equally capable of halting harmful invading phage DNA as they are halting beneficial mobile genetic elements, including those encoding antibiotic resistance and pathogenicity genes (7–9). As such, some pathogens only have alternative anti-phage defense systems (10). For example, the currently circulating biotype of epidemic *Vibrio cholerae*, the causative agent of the diarrheal disease cholera, does not rely on CRISPR-Cas for phage defense (11). Instead, *V. cholerae* evolved to use phage inducible chromosomal island-like elements (PLEs) to defend against the prevalent lytic phage, ICP1 (12). PLEs are mobile genetic elements that reside integrated in the small chromosome (chromosome II) of *V. cholerae* (12). During ICP1 infection of PLE(+) *V. cholerae*, PLE excises from the host chromosome, replicates to high copy and is horizontally transduced to naïve neighboring cells, all the while inhibiting phage replication through unknown mechanisms (Fig. 1).

**Figure 1.**
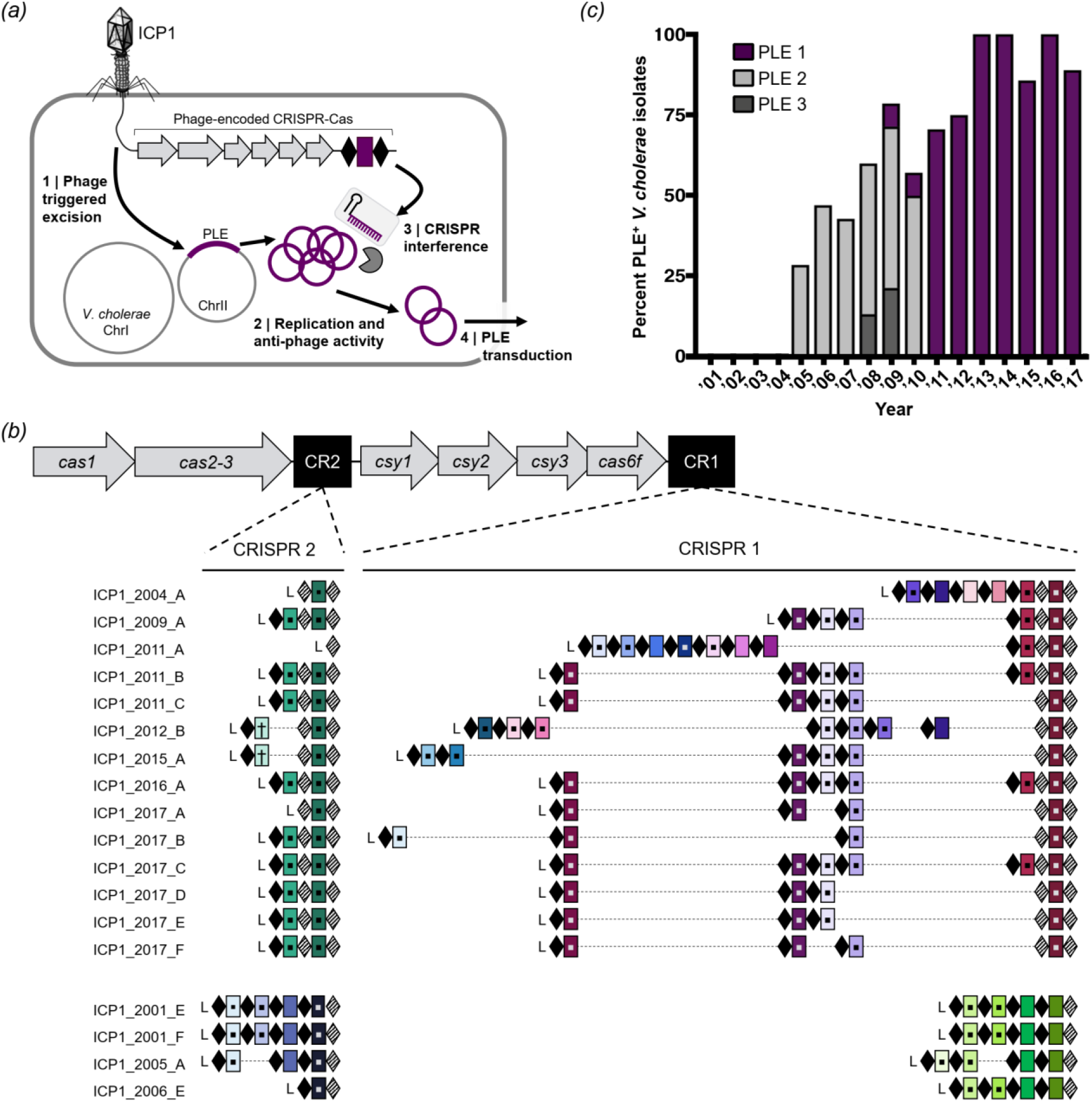
ICP1 uses CRISPR-Cas to overcome epidemic *V. cholerae* PLE. *(a)* Lytic phage ICP1 infects *V. cholerae* triggering PLE excision (20). PLE replicates and exerts anti-phage activity, ultimately leading to PLE transduction. Concurrently, ICP1-encoded CRISPR-Cas is expressed to interfere with PLE activity. *(b)* The architecture of the ICP1 CRISPR-Cas system and comparison of spacer composition between phage isolates. For each CRISPR locus, the repeat (28 bp) and spacer (32 bp) content is detailed as black diamonds and colored rectangles, respectively. Repeats that match the repeat consensus (13) are shown in solid diamonds, and degenerate repeats are indicated in hatched black diamonds. An AT-rich leader sequence (L) precedes each CRISPR locus. Identical spacers shared between isolates are shown as rectangles with identical colors. Spacers containing a square target PLE, and spacers containing a cross target the *V. cholerae* large chromosome. *(c)*Percentage of *V. cholerae* isolates harboring PLE recovered from epidemic sampling at the ICDDR,B over time (n=230 strains analyzed).

In order to overcome the anti-phage activity encoded by *V. cholerae* PLE, some ICP1 isolates use a Type I-F CRISPR-Cas system that directly targets PLE (Fig. 1), making the CRISPR-Cas system essential for the phage to form plaques on PLE(+) *V. cholerae* (13). Type I-F systems are composed of three Csy proteins that make up the Csy complex along with Cas6f, a protein involved in crRNA processing (14). This complex interacts with the processed crRNA to search DNA for a complementary protospacer with an appropriate self versus non-self discrimination sequence, known as the protospacer adjacent motif (PAM) (15). Upon finding a match with an appropriate PAM, the trans-acting Cas2-3 fusion protein is recruited to degrade the target DNA. In addition to exonuclease activity, Cas2-3 is a processive helicase *in vitro*, allowing for continued degradation of the target DNA (16,17). Recently, sequence analysis identified phages that are predicted to encode CRISPR arrays and/or Cas genes (18,19); however, ICP1 is the only phage shown to encode a fully functional CRISPR-Cas system (12,13).

As is true when CRISPR-Cas is harnessed by a prokaryotic host for genome defense, the ICP1-encoded CRISPR-Cas system is tasked with targeting and degrading a hostile mobile genetic element. However, there are additional challenges associated with a phage encoding and relying on CRISPR-Cas for its own survival. The ICP1 infection cycle occurs over a 20 minute period, and current data suggest that ICP1 synthesizes its CRISPR-Cas machinery *de novo* upon infection of *V. cholerae* (13). PLE is induced to excise within minutes of infection through interactions with an early phage-encoded gene product (20). Thus, in order to overcome PLE, CRISPR synthesis and interference must outpace a rapidly replicating target.

ICP1 and *V. cholerae* are consistently co-isolated from patient stool samples in regions where cholera is endemic such as Bangladesh (12,21,22). Five genetically distinct PLE variants in *V. cholerae* have appeared in temporally discrete waves across cholera epidemics (12). Previous analysis revealed that ICP1-encoded CRISPR-Cas can adapt and acquire new spacers against PLE under laboratory conditions (13), however the rules governing spacer acquisition and targeting efficacy for this system are not known. Further, recent comparative genomics of 18 ICP1 isolates collected from Bangladesh between 2001-2012 found that 50% carry CRISPR-Cas (23), however the contemporary state of circulating ICP1 and *V. cholerae* PLE in the region are not known.

Here, we provide an up-to-date understanding of the genomic variants of ICP1 and PLE circulating in Bangladesh. We find that natural ICP1 isolates encode multiple anti-PLE spacers and experimentally validate that increased PLE targeting by ICP1 is required to fully abolish PLE mobilization. Significantly, using a high-throughput spacer acquisition assay and experimental validation, we show that noncanonical PAMs and indirect protospacers in the *V. cholerae* small chromosome can unexpectedly provide protection against PLE. Our results support a model in which ICP1-encoded CRISPR-Cas that is directed against the *V. cholerae* small chromosome is in a race to reach PLE before it excises from the chromosome to exert its anti-phage activity. Taken together, our study highlights the differences between interference competent spacers under laboratory conditions and those that are selected for in nature to provide mechanistic insight into the evolutionary pressures governing the interactions between epidemic *V. cholerae* and its longstanding battle with the predatory phage ICP1.

## Methods

### Strains, growth conditions and genomic analysis

Phage, bacterial strains and plasmids used in this study are listed in Supplementary Tables 1-3. Bacteria were routinely grown at 37 °C on lysogeny broth (LB) agar or in LB broth with aeration. Media was supplemented with ampicillin (50 μg/ml), kanamycin (75 μg/ml), spectinomycin (100 μg/ml), and/or streptomycin (100 μg/ml) when appropriate. Phage susceptibility was determined by standard soft agar overlays as described (11) and phage plaque spot plates were performed as described previously (20). Cholera stool samples collected and stored at the ICDDR,B between 2015-2017 were probed for the presence of phage by standard soft agar overlays, and *V. cholerae* isolates were recovered by plating on Thiosulfate Citrate Bile Salts Sucrose selective media (Difco). ICP1 specific primers (13,22) and PLE specific primers (Supplementary Table 4) were used for preliminary screening of isolates from stool samples. The presence of CRISPR-Cas in ICP1 and PLE in *V. cholerae* was validated by whole genome sequencing. Genomic libraries were generated using NEBNext Ultra II DNA Library preparation kit for Illumina (New England Biolabs), according to the manufacturer’s recommended protocols. Paired-end sequencing (2 × 150 bp) was performed on an Illumina HiSeq4000 (University of California, Berkeley QB3 Core Facility). Sequencing assembly/mapping and detection of CRISPR was performed as described (23). The *V. cholerae* clinical isolate KDS1 genome was sequenced on Illumina HiSeq4000, PacBio Sequel and Oxford Nanopore MinION sequencers (University of California, Berkeley QB3 Core Facility). Assembly of KDS1 sequences was performed using the canu assembler v1.6 (24) to combine the PacBio and Oxford Nanopore reads into genomic scaffolds for the large and small chromosomes using default settings and an expected genome size of 4033460bp. This generated two scaffolds of the expected sizes for each chromosome which were then polished with the Illumina paired-end sequences using Pilon v1.22 (25) with the “fix all” command to generate a high-quality genomic assembly in a fasta format of both chromosomes (Supplementary File 1).

*V. cholerae* mutants were constructed by natural transformation as described (26). Mutations in ICP1 were generated using CRISPR-Cas mediated genome engineering with the *V. cholerae* classical biotype Type I-E system as described (11) (Supplementary Table 3). Engineered phage +/- Cas1 D244A with a spacer 9 were validated by plaquing on a permissive PLE 1 host and determining the frequency of phage with a newly acquired spacer by calculating the efficiency of plaquing on the permissive PLE 1 host to a PLE 1 host with the protospacer deleted. Examination of PLE replication and transduction during phage infection was described as reported previously (12).

### High throughput spacer acquisition, data processing and analyses

Three independent experiments were performed as follows: A 50 mL culture of PLE 1 *V. cholerae* was grown to OD_600_ = 0.3 and infected with ICP1_2011_A ΔS9 (13) at an MOI of 1. Infected cells were incubated for 90 minutes at 37 °C with aeration, at which point lysis was observed. The lysate was treated with chloroform and centrifuged to remove bacterial debris. Phage were precipitated with 10% (w/v) polyethylene glycol (PEG) 8000 at 4°C overnight. Phage pellets were collected by centrifugation at 4°C and the passaging was repeated as above. After three passages, the resulting pools were plated on the PLE 1^S8*^ host to select for phage with expanded arrays that allow plaque formation. Phage DNA libraries were generated by homopolymer tail-mediated PCR (HTM-PCR) as previously described (27). As ICP1_2011_A possesses only a single functional CRISPR array (Fig. 1b), the expanded phage CRISPR 1 array was amplified from genomic DNA libraries by PCR using custom barcoded primers (Supplementary Table 4) to sequence the leader proximal spacer. 50bp single-end sequencing was performed on an Illumina HiSeq 2500 (Tufts University Core Facility) using a custom sequencing primer. The resulting reads as fastq files (Supplementary File 2) were mapped to the large and small chromosome of *V. cholerae* strain KDS1 using Bowtie v1.2.2 (28) with a seed_length of 31 and allowing for 0 max_total_mismatches which ensured that spacer to protospacer matches were 100% identical. These mappings were performed in two parallel ways: first, to obtain all possible spacer mapping locations regardless of the number of identical protospacer targets (i.e. translucent spacers in Fig 3b) and second, restricting max_alignments to 1 which only mapped spacers with exactly one unique mapping location across both chromosomes. With a custom Python script (https://git.io/fNVqZ) we extracted the PAM sequences and GG PAM slippage locations from the restricted unique mappings. We also used this script to generate spacer mapping location graphs for both set of mappings using Biopython’s GenomeDiagram module (29).

## Results

### ICP1-encoded CRISPR-Cas is fixed in the natural phage population

We set out to compare ICP1 and PLE from contemporary cholera patient stool samples to previously identified isolates from the International Centre for Diarrhoeal Disease Research, Bangladesh (ICDDR,B) in Dhaka, Bangladesh (13,21). We isolated eight new ICP1 isolates from cholera patient stool samples collected between 2015-2017 and found that all isolates harbor CRISPR-Cas. Thus it appears that ICP1 isolates lacking CRISPR have not been identified in Bangladesh since 2006 (23). Analysis of the CRISPR arrays indicates a strong selection for spacers specifically targeting PLE (Fig. 1b, Supplementary Table 5), supporting the function of the ICP1-encoded CRISPR-Cas system as a counter-attack against the anti-phage island PLE (13). To evaluate if the fixation of CRISPR in ICP1 is necessitated by co-circulating PLE in epidemic *V. cholerae*, we determined the prevalence of PLE over the same near two-decade long period in Dhaka. Combined with previous analyses (12,21), we observed an increase in the prevalence of PLE(+) *V. cholerae* in epidemic sampling over time (Fig. 1c). Of note is the high prevalence of PLE 1 *V. cholerae* over the past 6 years, indicating that this variant of the anti-phage island is currently dominating the epidemic landscape in Dhaka. Despite the relatively long period over which PLE 1 has been dominant in Dhaka, and consistent with previous results (12,21), whole genome sequencing of eight PLE 1 *V. cholerae* isolates showed that PLE 1 is 100% identical at the nucleotide level in all strains.

### Multiple spacers increase ICP1 CRISPR-Cas mediated PLE interference

All of the natural phage we isolated encode multiple CRISPR spacers against PLE (Fig. 1b); however, previous work revealed that only one functional spacer is required for ICP1 to overcome PLE-mediated anti-phage activity as evaluated by plaque formation (13). Conversely, a single spacer against the PLE did not prevent transduction of PLE (12). To investigate the consequences of varying spacer number and identity on PLE transduction and replication, we used co-isolated ICP1 and PLE 1 *V. cholerae* obtained from a cholera patient sample in 2011 (13). This ICP1 isolate harbors two spacers (spacers 8 and 9) at the leading edge of the CRISPR 1 array that target PLE 1. We also used an isogenic phage with a spontaneous loss of spacer 9 (13), as well as one that acquired an additional 10^th^ spacer targeting PLE *in vitro* (Fig. 2a). Despite the ability to overcome PLE and form plaques, spacer 8 targeting was not sufficient to decrease PLE transduction during ICP1 infection relative to an untargeted control (Fig. 2b). In comparison, two anti-PLE spacers decreased PLE transduction during ICP1 infection and three spacers completely abolished PLE transduction, showing that increased CRISPR targeting by ICP1 has a stronger anti-PLE effect. To evaluate potential differences between spacer 8 and spacer 9 on PLE targeting, we used PLE 1 with a protospacer mutation (PLE 1^PS8^*) that inhibits spacer 8-mediated PLE targeting (13). Strikingly, just spacer 9 targeting PLE alone was able to decrease PLE transduction to the same level as when two spacers were targeting PLE (Fig. 2b).

**Figure 2.**
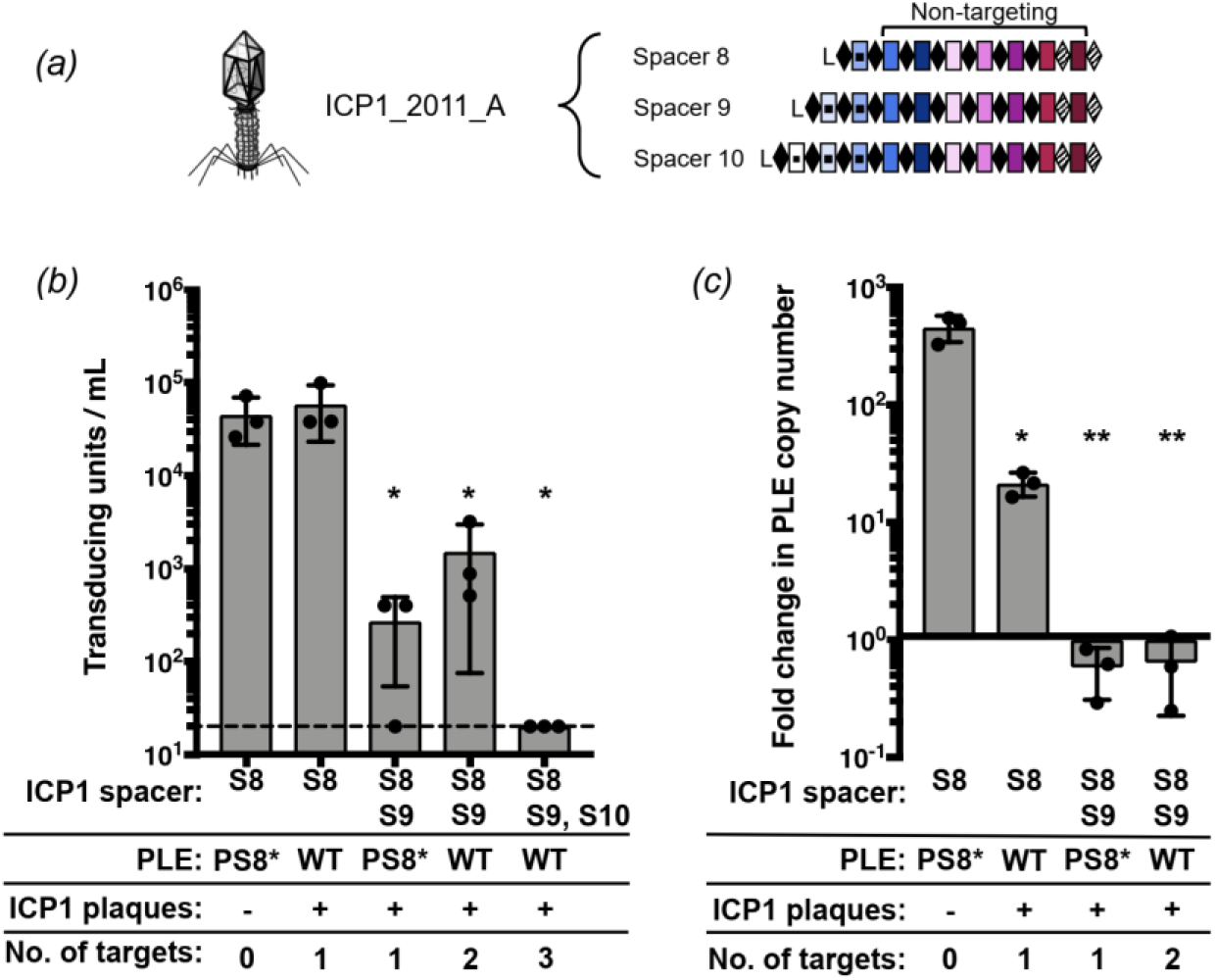
CRISPR can limit horizontal transmission of PLE. *(a)* ICP1_2011_A with anti-PLE spacers S8, S9 and S10 tested in panels b and c. *(b)* PLE transduction after infection with ICP1 with 0,1,2 or 3 spacers. The dashed line indicates the limit of detection for this assay. A single spacer is necessary and sufficient to permit lytic growth of ICP1 on PLE 1 *V. cholerae* as seen by equal plaque formation. *(c)* PLE replication 20 minutes after infection with ICP1 with 0,1 or 2 spacers as determined by qPCR. For panels b and c, error bars indicate standard deviations of biological triplicates. Significance was determined by T Test, * p<0.05, ** p<0.005.

We next analyzed the copy number of PLE during infection with the ICP1 encoding one or two targeting spacers to identify if the differences in reducing PLE transduction were due to differences in PLE copy number (Fig 2c). In the absence of ICP1 CRISPR targeting, PLE replicates to high copy number, which facilitates horizontal transmission. Targeting with only one spacer was sufficient to significantly decrease PLE replication, and in agreement with the transduction data, spacer 9 had a stronger inhibitory effect on PLE replication than spacer 8. Altogether, these results demonstrate that not all spacers selected in nature equally interfere with PLE mobilization and that increasing the number of spacers provides enhanced capacity of ICP1 to interfere with PLE.

### Interference-driven spacer acquisition in ICP1 reveals indirect targets and non-canonical PAMs

Since spacer composition variability in nature was lower than we expected (Fig 1b), we next set out to experimentally sample the repertoire of spacers that ICP1 can acquire to overcome PLE. Low-throughput experiments previously demonstrated that ICP1 can acquire new spacers targeting the PLE under laboratory conditions without the need to overexpress *cas* genes (13). To further analyze the natural process of interference-driven spacer acquisition in this system, we performed high-throughput sequencing of expanded CRISPR arrays of phage selected on PLE 1 *V. cholerae*. We infected PLE 1 *V. cholerae* with ICP1 containing spacer 8 (Fig 2a), and the recovered lysate was probed for ICP1 progeny with newly acquired spacers that allowed for plaque formation on a PLE 1^PS8*^ host. Illumina sequencing of the leader-proximal spacer in CRISPR 1 allowed us to sample over 10^6^ acquired spacers in each replicate experiment (Supplementary Table 6). In order to accurately map the spacers to the PLE 1 *V. cholerae* host, we performed complete whole-genome sequencing and assembly of the bacterial genome (Supplementary File 1). As was previously reported (12), we found that PLE 1 was integrated in a *V. cholerae* repeat (VCR), of which over 100 repeats intersperse the *V. cholerae* small chromosome in a gene-capture region, the superintegron (30). In total, 96% of the acquired spacers mapped to PLE, while, interestingly, the other 4% mapped to *V. cholerae* chromosomes (Supplementary Table 6).

Mapping of the spacers to the small chromosome showed a pattern of strand bias that reflected previous observations in primed acquisition experiments performed in other Type I-F systems (31), with a distribution of acquired spacers 5’ of the protospacer on the non-targeted strand and 3’ of the protospacer on the targeted strand (Fig. 3a, Supplementary Fig. 2). The distribution of spacers acquired 5’ of the protospacer on the nontargeted were split between the small chromosomal region proximal to the PLE 1 integration site (Fig. 3b), as well as the 3’ end of PLE. Acquired spacers mapping to the *V. cholerae* chromosome were not evenly distributed between the large and small chromosome, but instead ~90% of the chromosomal spacers mapped to the small chromosome (Supplementary Table 6, Fig. 3b). Spacers that mapped to the large chromosome were restricted to a mu-like region (Fig. 3c), which was duplicated in this strain and was also in the small chromosome proximal to PLE (Fig. 3b). Acquired spacers mapped uniformly throughout the superintegron, however, this is likely an artifact as the superintegron is highly repetitive. When considering spacers that map to a single site in the small chromosome, we observed an obvious bias for acquired spacers mapping closer to the PLE integration site (Fig. 3b).

**Figure 3.**
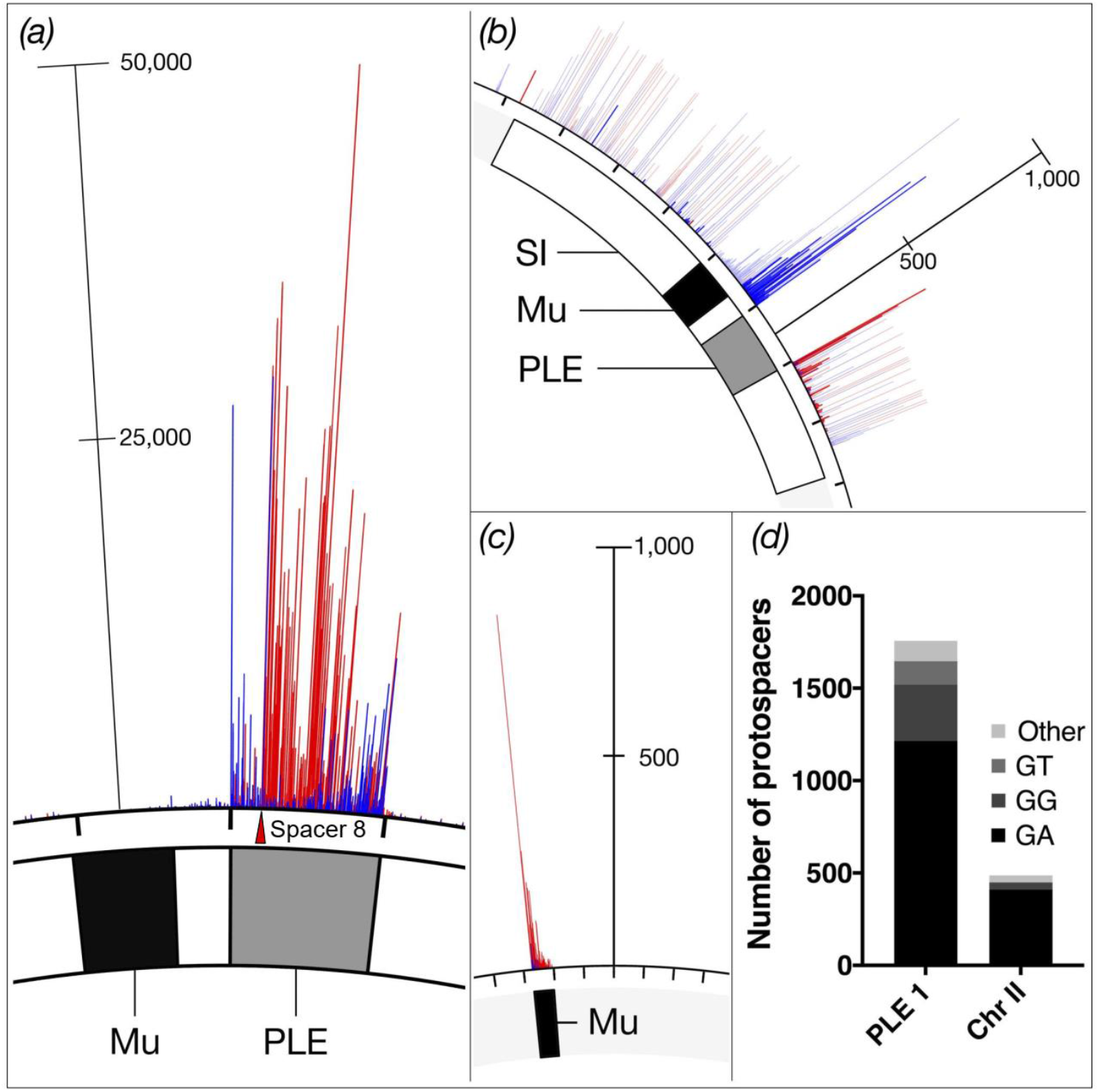
High-throughput interference driven spacer acquisition mapping. *(a)* The locations of the ICP1 CRISPR leader-proximal spacer on *V. cholerae* small chromosome. The location of the interference-efficient spacer (S8) is indicated with the red triangle. *(b)* Spacer locations on the *V. cholerae* small chromosome (PLE mappings not shown for clarity). Uniquely mapped spacers are shown in solid blue or red, while translucent bars show mapping of spacers to all possible locations. *(c)* Spacer locations on the large chromosome. For panels a, b and c, spacers on the plus and minus strand are indicated in red and blue, respectively. The scale bar measures the number of mapped spacers, and the tick marks around the chromosome are in 18kb intervals. The white box represents the superintegron (SI), the black box is the mu-like region and the grey box is PLE 1. *(d)* Proportion of unique protospacers with a GA or other dinucleotide PAM sequence in PLE or in the small chromosome.

Consistent with CRISPR+ ICP1 isolates from nature (Supplementary Figure 1), the majority (~70%) of the spacers acquired experimentally targeted protospacers in PLE 1 that were flanked by a 3’ GA PAM (Fig. 3d). However, ~30% of protospacers in PLE had non-canonical PAMs, and of those, the majority were GG or GT. Previous CRISPR acquisition studies in Type I-F systems indicate that alternative PAMs can be explained by a “slippage” event (31,32). To identify putative slippage events, we analyzed the sequences adjacent to GG PAMs and found that 45% of GG PAMs have a canonical GA within 3 nucleotides of the PAM position, suggesting that the ICP1 acquisition machinery has a propensity to slip (Fig. 4a).

**Figure 4.**
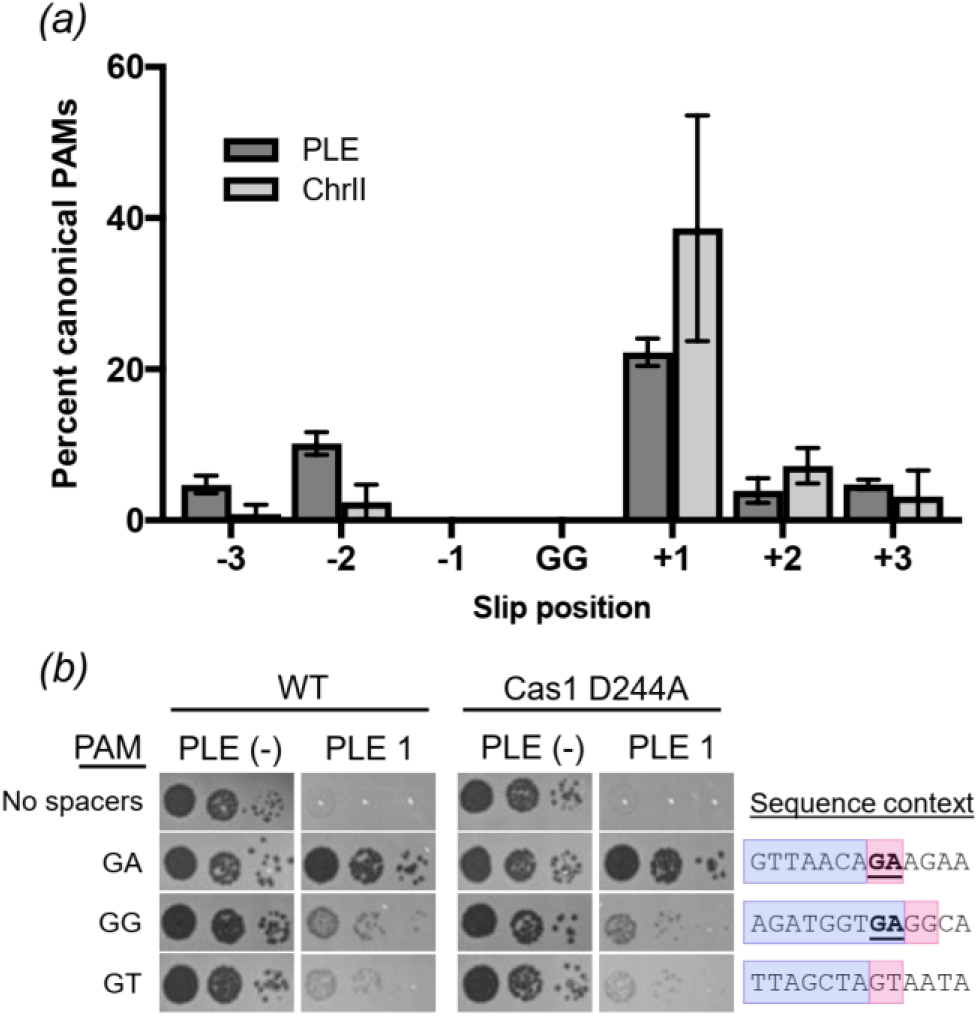
Characterizing non-canonical PAMs. *(a)* Frequency of canonical GA PAM +/- 3nt from non-canonical GG PAM across all data sets. *(b)* Tenfold dilutions of ICP1 engineered to contain a spacer that targets PLE 1 with a non-canonical PAM spotted on *V. cholerae* PLE(-) or PLE 1 lawns showing the ability of different phage strains to form plaques (dark spots, zones of killing) (left). Sequence context (right) of the region adjacent to the PAM. The protospacer is boxed in purple and PAM is boxed in pink. The consensus canonical PAM GA is bolded and underlined.

We next wanted to determine if these non-canonical PAMs are functional for PLE interference. To do so, we engineered ICP1 to encode a single spacer reflective of an experimentally acquired spacer with the most common non-canonical PAMs: either a GG or GT (Fig. 3d) and evaluated plaque formation on PLE 1 *V. cholerae*. Despite relying on a non-canonical PAM, we found that ICP1 is able to target those protospacers and overcome PLE, albeit at a lower efficiency than when targeting a protospacer with a canonical GA PAM (Fig. 4b). Even when no canonical PAM was within +/- 3 nt, ICP1 was still able to overcome PLE targeting a protospacer with a GT PAM. As PAM mutations are frequently a source for primed acquisition (33), we tested if the observed residual CRISPR activity was due to further spacer acquisition and interference. We constructed a Cas1 D244A mutation, which disrupts a conserved metal coordinating residue to inhibit spacer acquisition (32) (Supplementary Fig. 3) and tested if plaque formation was altered (Fig 4b). We observed no difference in the efficiency of plaque formation between the Cas1 mutants and the parental phage, suggesting that the ICP1 CRISPR-Cas system is more tolerant of divergent PAMs during infection than previously characterized (13).

### Protospacers in the small chromosome facilitate ICP1 CRISPR-Cas-mediated PLE interference

In our spacer acquisition experiment we identified a subset of spacers that target a mu-like region in the *V. cholerae* large chromosome (Fig. 3c), suggesting that CRISPR targeting of the mu-like region was advantageous in overcoming PLE. To test the role of the mu-like region protospacer in PLE interference, we isolated ICP1 that had acquired a spacer that targets the mu-like region and was able to form plaques on PLE 1^PS8^* (Fig 5a). Since assembly of the *V. cholerae* genome revealed that the mu-like region was present in each chromosome (Fig 5a) we wanted to evaluate if targeting the mu-like region *per se* was allowing for plaque formation, or if the chromosomal context was important in allowing for CRISPR-meditated interference with PLE. To test this difference, we generated a single knockout of the mu-like region in the large chromosome and a double knockout in both chromosomes. ICP1 CRISPR-mediated interference with PLE was abolished in the double knockout, however, knocking out the mu-like region in the large chromosome had no effect on ICP1 plaque formation (Fig 5b). These results show that CRISPR targeting of the *V. cholerae* large chromosome is dispensable for phage overcoming PLE, while targeting the small chromosome is sufficient to overcome PLE activity.

**Figure 5.**
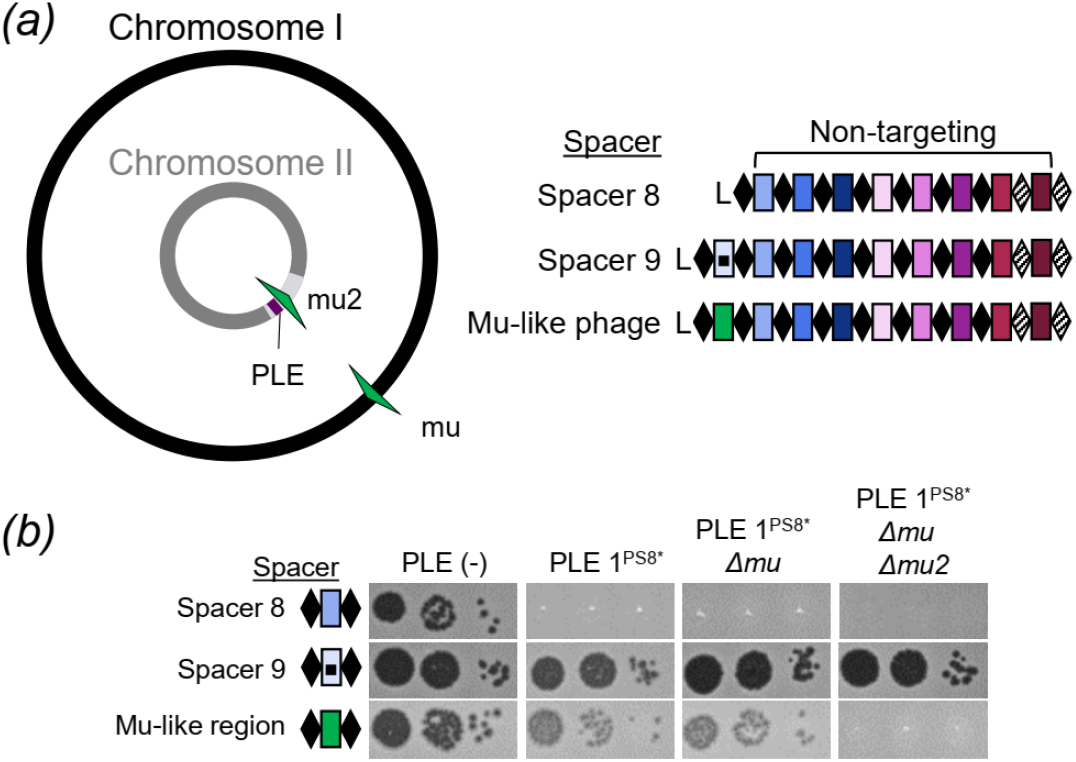
ICP1 CRISPR-targeting of the small chromosome facilitates PLE interference. *(a)* Cartoon (left) of the *V. cholerae* large and small chromosomes. The superintegron is shown in light grey, the PLE is shown in purple. The two mu-like regions in the large and small chromosome are shown in green arrows. ICP1_2011_A CRISPR variants (right) used to test the role of targeting sites. *(b)* Tenfold dilutions of ICP1 with the spacers indicated spotted on *V. cholerae* lawns showing the ability of different phage strains to form plaques.

### When CRISPR goes off target: going the distance to maintain interference

As processivity of Cas2-3 has been demonstrated *in vitro* (17), we speculated that ICP1 targeting of the small chromosome proximal to PLE interferes with PLE anti-phage activity by the processive degradation of PLE along with the chromosome; however, PLE excises from the chromosome early during ICP1 infection (20). This timing suggests that CRISPR targeting and Cas2-3 processive degradation of the small chromosome would have to happen prior to PLE excision and would therefore likely be distance dependent. In support of this hypothesis, experimentally acquired spacers mapping to the small chromosomal clustered proximal to PLE (Fig. 3b). To test the impact of targeting at increasing distances from PLE, we engineered ICP1 to possess CRISPR arrays containing only one spacer drawn from the experimental acquisition pool that targets the small chromosome at varying distances away from PLE. We then assayed the ability of these engineered phage to overcome PLE and form plaques (Fig. 6a). As a positive control, ICP1 engineered with a spacer that targets internal to PLE formed robust and equal plaques on PLE(-) and PLE 1 hosts. In comparison, phage with a spacer that targets far (>400 kb) from PLE were unable to form plaques on PLE 1. Conversely, ICP1 that target a protospacer only 0.5, 1.5 or 2.5 kb from PLE were able to efficiently overcome PLE and form plaques. Phage targeting protospacers at intermediate distances away from PLE (>20 kb) demonstrated weak plaque formation on PLE 1. Surprisingly, we observed that ICP1 with some spacers targeting relatively far from PLE (53 and 46kb away) were still able to form robust plaques on PLE 1 (Fig 6a). While all of the spacers selected for this assay had one perfect protospacer match in the chromosome (and have a GA PAM), we identified >100 putative promiscuous target sites for these spacers which would bring the chromosomal target much closer to PLE 1, which may explain these phage’s ability to overcome PLE. To test if spacer acquisition had a role in plaque formation, we engineered Cas1 deficient ICP1 in each CRISPR proficient chromosomal targeting phage and assayed for plaque formation on the PLE 1 host. Despite being unable to acquire spacers (Supplementary Fig. 3), the phage retained the same plaquing phenotype. We quantified the weaker plaque formation observed when ICP1 targets >20 kb away from PLE 1 by measuring plaque size compared to PLE (-) *V. cholerae* (Fig 6b). As compared to phage with PLE internal and PLE proximal spacers, phage with chromosomal spacers targeting >20 kb away from PLE had significantly limited plaque size. These results indicate that some PLE-mediated anti-phage activity is retained when CRISPR-Cas is directed at increasing distances from PLE in the small chromosome.

**Figure 6.**
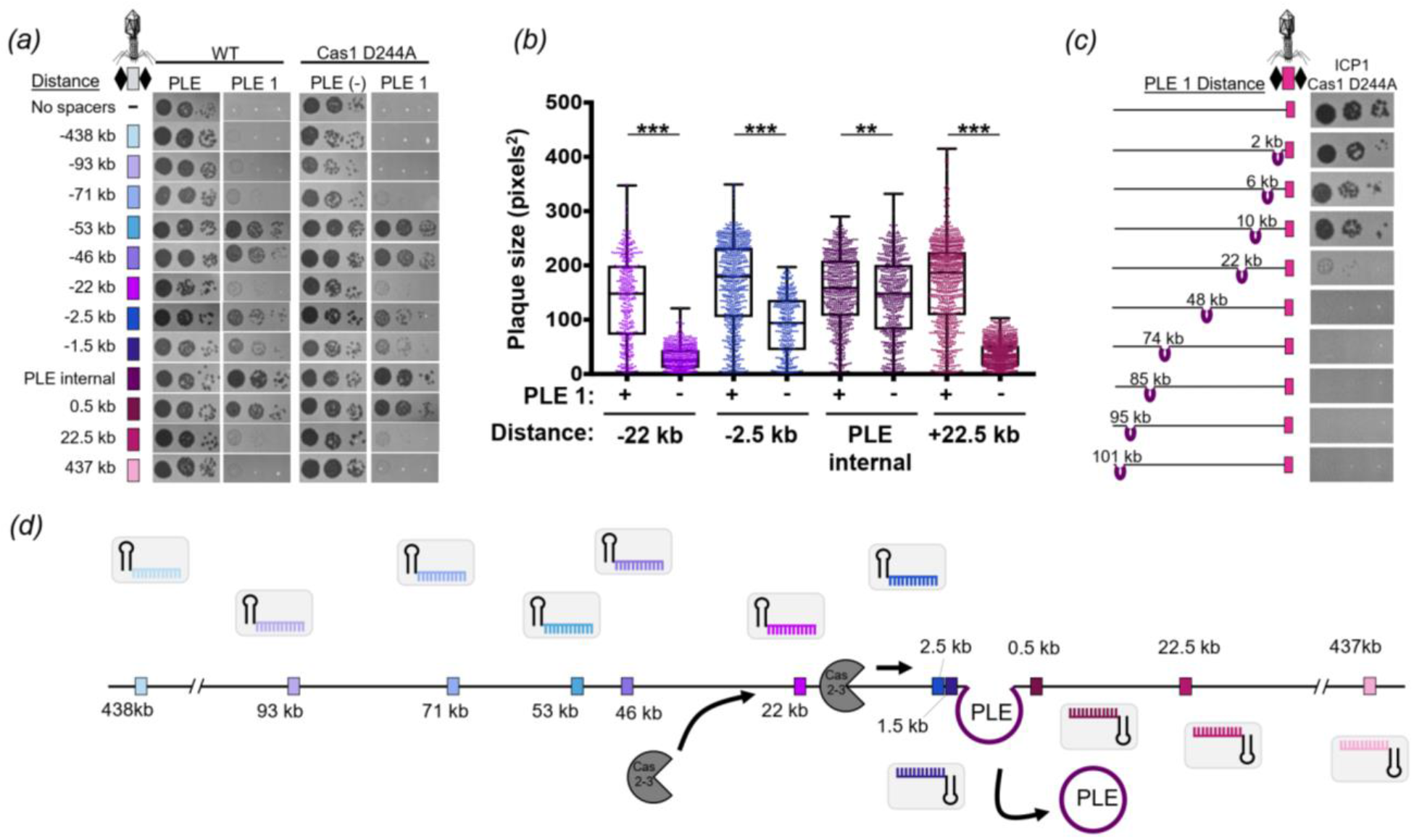
Interference potential of spacers directed to the small chromosome is dependent on proximity to PLE. *(a)* Tenfold dilutions ICP1 engineered with a chromosomal protospacer +/- Cas1 spotted on lawns of *V. cholerae. (b)* Plaque size of ICP1 variants plated with *V. cholerae*. The distance of the chromosomal protospacer from PLE 1 integration site is indicated. Significance was determined by Mann-Whitney U Test, **p<0.005, ***p<0.0001. *(c)* Tenfold dilutions of ICP1 engineered with a chromosomal protospacer spotted on lawns of *V. cholerae* harboring PLE in different locations in the chromosome. *(d)* Model of race between ICP1 Cas2-3 processive degradation of the *V. cholerae* chromosome and ICP1-mediated PLE excision. Csy complexes (grey boxes) with crRNAs (colored) search for a complementary protospacer (colored rectangles, experimentally assessed in panel a). Cas2-3 (dark grey) is recruited to the protospacer and processively degrades the DNA towards PLE (purple). ICP1 is able to form plaques when Cas2-3 degrades PLE before PLE excises from the chromosome, which occurs within 5 minutes of ICP1 infection.

To control for differences in spacer sequences, we also varied the location of the PLE and tested the ability of ICP1 with a single chromosomal spacer targeting the small chromosome to interfere with PLE 1. Following ICP1-mediated transduction, PLE 1 integrates into the *V. cholerae* repeat (VCR) of the new host (12). We collected a pool of PLE 1 transductants where PLE was integrated at varying distances from the chromosomal protospacer and challenged these strains with ICP1. As a control, we determined that all of the tested PLE 1 *V. cholerae* hosts were susceptible to ICP1 CRISPR-Cas interference when ICP1 possessed a PLE internal spacer (Supplementary Fig. 4); consistent with our earlier finding, PLE integrated at an increasing distance away from the protospacer was less susceptible to ICP1-encoded CRISPR interference (Fig. 6c).

## Discussion

Our results reveal that the latest front in the ongoing arms race between contemporary isolates of epidemic *V. cholerae* and its predator ICP1 necessitate the persistence of the ICP1-encoded Type I-F CRISPR-Cas system to counter PLE-mediated anti-phage activity (Fig 1). By using a high-throughput spacer acquisition assay, we gained insight into the full range of spacers that can combat PLE. Interestingly, our experimental findings on acquisition and interference do not reflect the rather limited diversity of spacers that ICP1 maintains against PLE in nature. These results highlight that not all spacers are equally proficient for interference, and that coupled analysis of these competing mobile genetic elements from nature reveals the evolutionary benefits of a particular complement of spacers more so than laboratory-based studies. Despite a lack of clear evidence indicating where the ICP1-encoded CRISPR-Cas system originated, it serves as a tractable model through which we can examine the biology of an endogenous Type I-F CRISPR-Cas system against its cognate foe.

Co-culture studies competing phage against CRISPR-Cas proficient bacterial hosts demonstrated that mutational escape by phage is limited by bacterial populations that have heterogenous CRISPR arrays (34). Here, we see that PLE 1 is highly conserved over time, even when co-circulating with CRISPR proficient ICP1. In light of previous suggestions, the diversity of CRISPR arrays in ICP1 populations may limit the success of PLE escape mutants. Surprisingly, however, we see very little diversity in the spacer composition of ICP1 CRISPR arrays with the same minimal spacers being conserved in phage circulating for over eight years (Fig. 1b). Likewise, CRISPR-proficient ICP1 isolated from nature always encoded more than one spacer against PLE, which would be expected to limit CRISPR escape mutations. It may be that there is limited room for genetic drift in the PLE genome, permitting ICP1 to streamline its CRISPR array, keeping only the most efficient spacers while also maintaining an advantageous genome size.

Akin to studies of bacterial Type I-F CRISPR-Cas mediated interference with plasmid transformation and conjugation (35), we similarly see that the spacer sequence and quantity of spacers in the array have a role in ICP1’s ability to abolish PLE spread (Fig. 2). This may be due to differences in crRNA abundance or stability, or sequence dependent subtleties that dictate interference potential, as has been proposed previously (36). Despite spacer 9’s improved interference with PLE mobilization compared to spacer 8, we still observed a slight defect in plaque size when comparing engineered phage with only spacer 9 relative to a PLE(-) host (Fig. 6b), suggesting that even this improved spacer alone is not sufficient to fully overcome PLE-mediated anti-phage activity. By encoding a seemingly redundant set of spacers targeting PLE, ICP1 increases its ability to overcome PLE and limit PLE spread in the environment.

As expected, the majority of spacers acquired in our high-throughput acquisition assay directly target PLE (Fig. 3a). Analysis of natural ICP1 isolates recovered from cholera patient stool samples shows that the phage-encoded CRISPR-Cas system recognizes a GA PAM, (Supplementary Fig. 1) which, although atypical for Type I-F systems (37), has been confirmed through single mutations to a C in both positions (13). Notably, we found that ICP1 was able to incorporate spacers that targeted non-canonical PAMs (Fig. 3d) and that these spacers can suffice for PLE interference (Fig. 4b). In comparison to another high throughput spacer acquisition assay in a Type I-F system, which found >90% of all protospacers flanked by the canonical PAM (31), it appears that the phage-encoded system is less discriminating with only 70% of protospacers flanked by the expected PAM. However, targeting a protospacer with a non-canonical PAM reduced the efficiency of plaquing compared to the canonical PAM (Fig 4b). As such, in nature ICP1 targeting a protospacer with a non-canonical PAM would not be able to completely interfere with PLE and thus would be selected against. This hypothesis is additionally supported by the observation that very few non-canonical PAM protospacers were associated with indirect targets in the small chromosome. As these chromosomal spacers are themselves less proficient for interference (Fig. 6a and 6b), the added disadvantage of targeting a protospacer with a non-canonical PAM likely tips the balance in favor of PLE, likely explaining the lower abundance of these spacers in our selection experiments.

Despite the presence of spacers that target the *V. cholerae* large chromosome in the high-throughput spacer acquisition assay (Fig. 3c), we show that targeting this chromosome is dispensable for CRISPR interference of PLE (Fig. 5b). Interestingly, two of the natural ICP1 isolates contain a spacer that targets a gene on the *V. cholerae* large chromosome (Fig. 1b). We speculate that this spacer was acquired from a *V. cholerae* strain possessing a duplication or rearrangement that is not represented in currently sequenced isolates, in which the protospacer was in the small chromosome proximal to PLE, allowing the phage to overcome PLE activity. However, this spacer does not seem to be maintained in the phage population, likely due to diminished PLE interference relative to PLE-direct spacers as we experimentally observed.

CRISPR targeting of the *V. cholerae* small chromosome can overcome PLE, but our results suggest a model in which there is a limit to the distance over which processive Cas2-3 degradation can occur to reach the PLE prior to excision (Fig. 6d), an action which occurs within five minutes of ICP1 infection that is directed by an early-expressed ICP1 protein (20). The limit of processivity appears to be around a distance of 23 kb (Fig. 6a and 6c), at which point either Cas2-3 is unable to continue to process along the *V. cholerae* chromosome or PLE excises before interference occurs. *In vitro* studies of Cas3 from Type I-E systems have demonstrated Cas3 translocation velocities of 89 to 300 bases per second and average processivities between 12 to 19 kb (38,39), however, the functional role and limitations of processivity *in vivo* are not known. Our results are the first indications of Cas2-3 processivity *in vivo*, with over 22 kb from a distal chromosomal protospacer over which CRISPR-Cas can maintain activity to overcome PLE. As this event must occur within five minutes of ICP1 initiating infection, the estimated processivity of ICP1 Cas2-3 is within the range of what has been reported for Type I-E Cas3, which is especially remarkable given the complexity of the crowded intracellular environment compared to simplified *in vitro* systems.

In comparison to other Cas nucleases like Cas9, which introduces a single double-stranded break (40,41), Cas2-3 degrades DNA as it translocates away from the protospacer (17), making it more likely to destroy and thus interfere with its target. This predicted advantage may account for the increased prevalence of Type I systems for phage defense (42). In the context of the battle between ICP1 and PLE, this processivity permits interference even with an indirect CRISPR target and has important implications for harnessing CRISPR-Cas in biotechnology and medicine. Since the characterization of the ICP1-encoded CRISPR-Cas system, phage engineered with CRISPR-Cas systems to target virulent, antibiotic resistant bacteria have been assayed for therapeutic applications (43,44), showing the value of innovating from natural systems to overcome disparate biological problems.

## Additional Information

### Acknowledgments

The authors are especially thankful to ICDDR,B hospital and laboratory staff for their support and would like to thank Shirajum Monira, Kazi Zillur Rahman, Fatema-tuz Johura, Marzia Sultana, and Monika Sultana in particular. The authors would also like to thank members of the Seed lab for critical feedback and thoughtful discussion regarding this manuscript.

### Ethics

The collection of cholera patient stools were approved by the ICDDR,B institutional review board. All samples were deidentified and written informed consent was obtained from adult participants and from the guardians of children.

### Data Accessibility

The datasets supporting this article have been uploaded as part of the Supplementary Material.

### Authors’ Contributions

ACM, KNL and KDS carried out the molecular lab work. AA developed and implemented tools for sequence data analysis. MA coordinated the collection of clinical specimens. ACM, MA and KDS conceived of the study. All authors participated in data analysis. ACM and KDS wrote the manuscript with input from all authors and all authors gave final approval for publication.

---

**Supplementary Figure 1.**
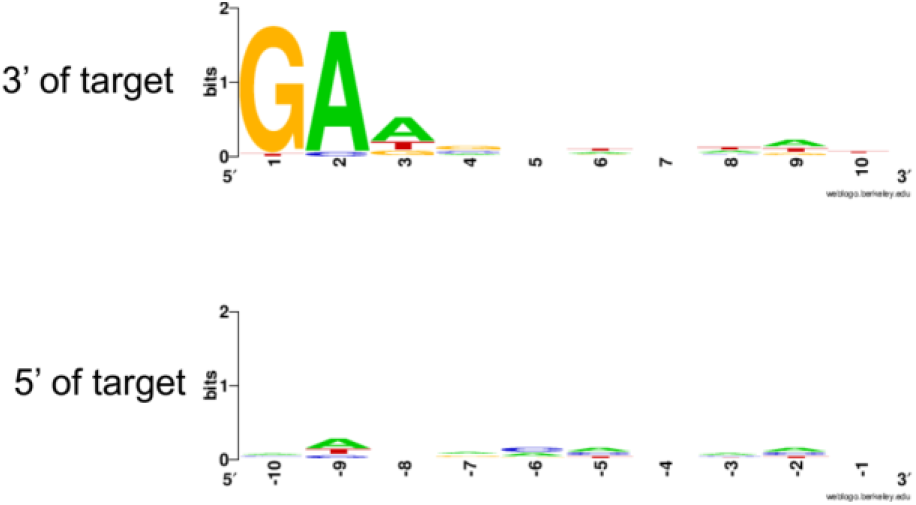
Sequence logo of PAMs of natural ICP1 isolates. The PAM sequence of the ICP1-encoded CRISPR-Cas system. Alignment of flanking sequence of all known targets of spacers found in natural ICP1 isolates. Sequence logos were generated using WebLogo (45).

**Supplementary Figure 2.**
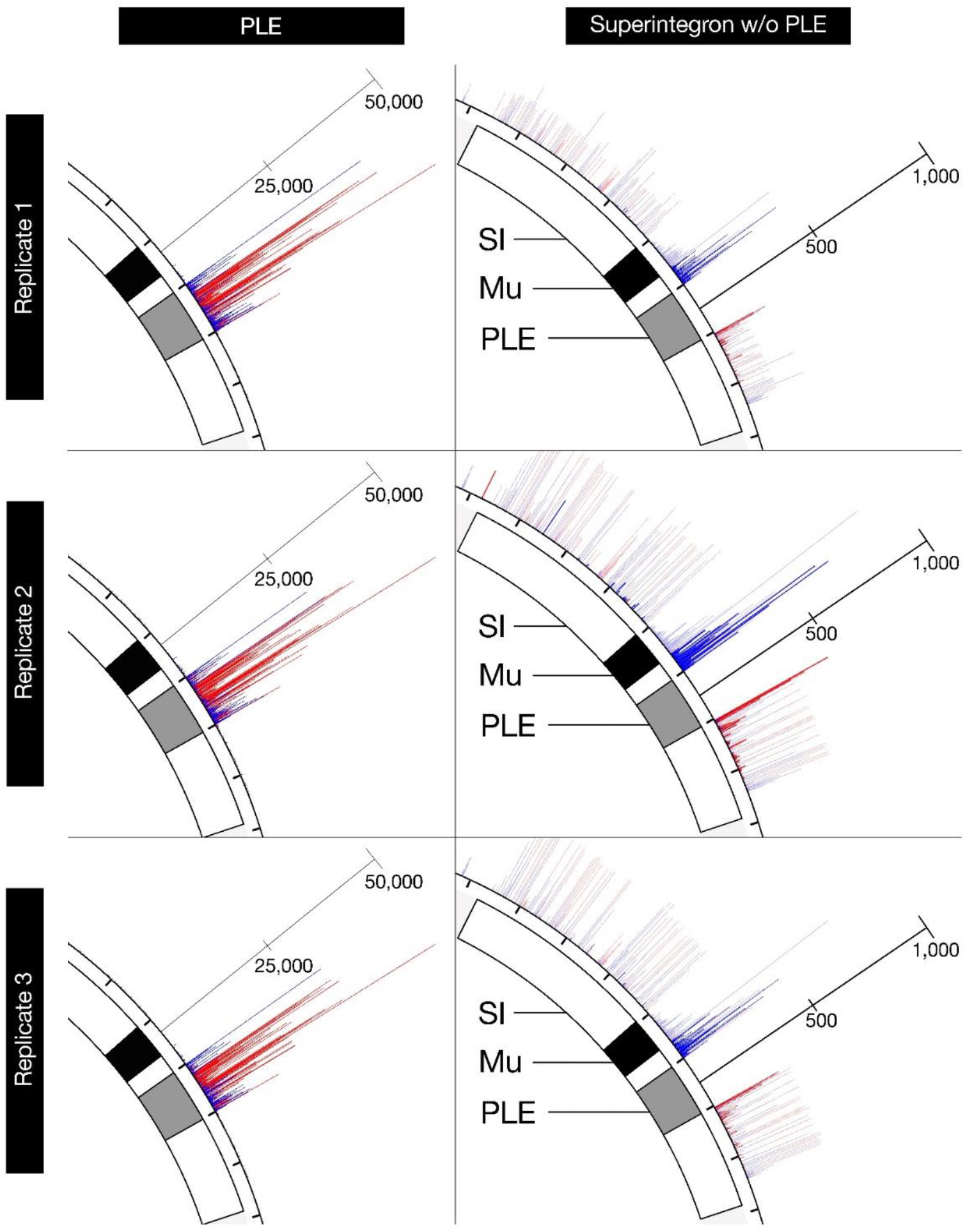
Replicates of high-throughput ICP1 spacer acquisition mapping. Spacer locations of the most leader-proximal spacers on the plus and minus strand are indicated in red and blue, respectively. Uniquely mapped spacers are shown in solid blue or red, while translucent bars show mapping of spacers to all possible locations. The scale bars measure the number of mapped spacers, and the tick marks around the chromosome are in 18kb intervals. Replicate 2 is the same data as in Figure 3a and b.

**Supplementary Figure 3.**
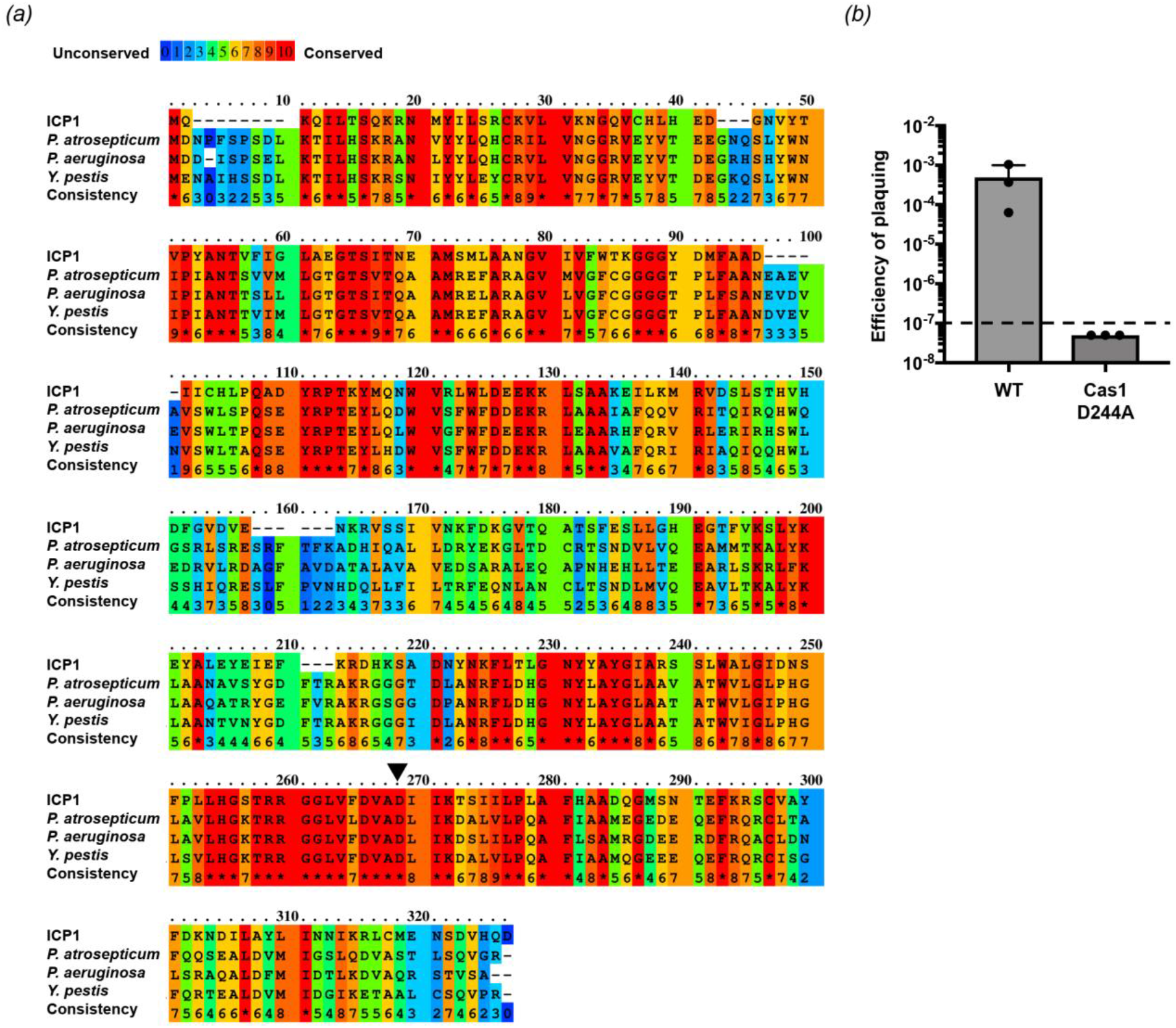
Conserved residue in Cas1 is necessary for spacer acquisition. *(a)* Praline alignment (46) of Cas1 from ICP1 and other Type I-F systems in the organism indicated. Black arrow indicates conserved residue mutated in ICP1. *(b)* Spacer acquisition was measured by the number of plaques on a PLE(+) host without a protospacer relative to the number of plaques on a PLE(+) host with a functional protospacer. Dashed line indicates level of detection. Error bars indicate standard deviation of three independent replicates.

**Supplementary Figure 4.**
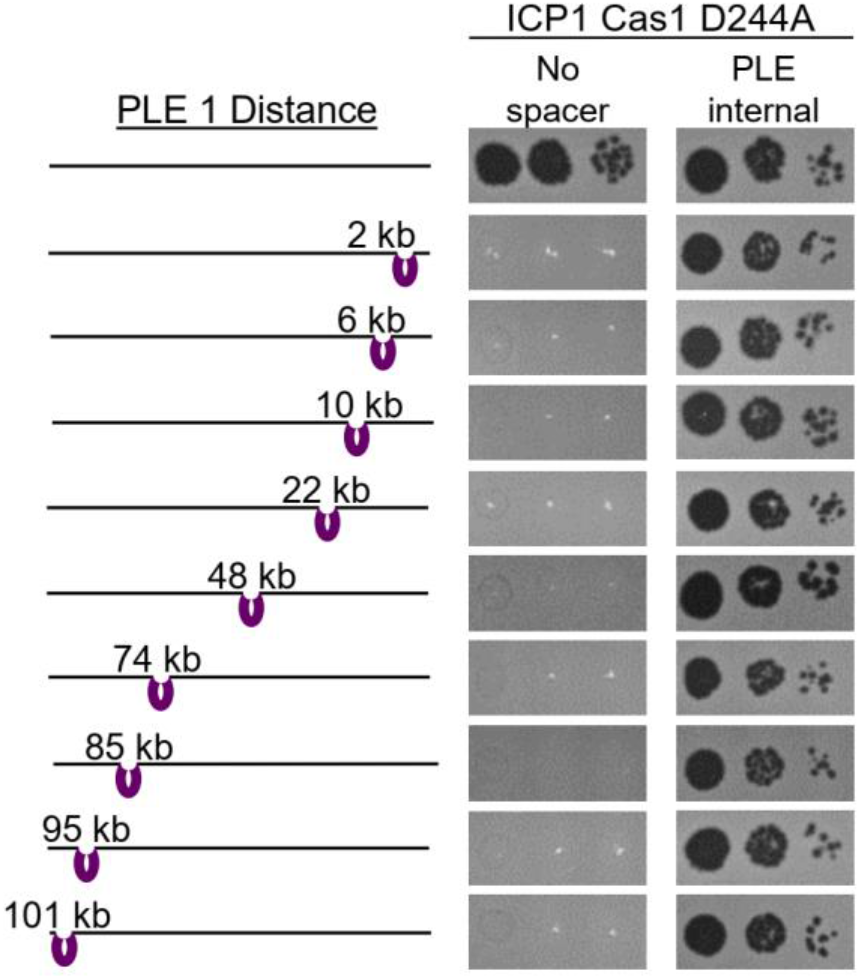
PLE transduced in unique sites in *V. cholerae* chromosome can be overcome by ICP1-CRISPR mediated interference. Tenfold dilutions ICP1 without any spacer or with a PLE targeting spacer spotted on lawns of *V. cholerae*. Transductants used are the same as in Figure 6c.

## References

1. Parikka KJ, Le Romancer M, Wauters N, Jacquet S. Deciphering the virus-to-prokaryote ratio (VPR): Insights into virus-host relationships in a variety of ecosystems. Biol Rev. 2017;92:1081–100.

2. Dy RL, Richter C, Salmond GPC, Fineran PC. Remarkable Mechanisms in Microbes to Resist Phage Infections. Annu Rev Virol. 2014;1(1):307–31.

3. Hille F, Richter H, Wong SP, Bratovič M, Ressel S, Charpentier E. The Biology of CRISPR-Cas: Backward and Forward. Cell. 2018;172(6):1239–59.

4. Barrangou R, Fremaux C, Deveau H, Richards M, Boyaval P, Moineau S, et al. CRISPR Provides Acquired Resistance Against Viruses in Prokaryotes. Science. 2007;315(5819):1709–12.

5. Sinkunas T, Gasiunas G, Fremaux C, Barrangou R, Horvath P, Siksnys V. Cas3 is a single-stranded DNA nuclease and ATP-dependent helicase in the CRISPR/Cas immune system. EMBO J. 2011;30(7):1335–42.

6. Koonin E V., Makarova KS. Mobile genetic elements and evolution of CRISPR-Cas systems: all the way there and back. Genome Biol Evol. 2017;9(10):2812–25.

7. Shmakov, Sergey; Sitnik, Vassilii; Makarova, Kira; Wolf, Yuri; Severinov, Konstantin; Koonin E. The CRISPR Spacer Space Is Dominated by Sequences from Species-Specific Mobilomes. MBio. 2017;8(5):e01397–17.

8. Palmer KL, Gilmore MS. Multidrug-Resistant Enterococci Lack CRISPR-Cas. MBio. 2010;1(4):e00227–10.

9. Bikard D, Hatoum-Aslan A, Mucida D, Marraffini LA. CRISPR interference can prevent natural transformation and virulence acquisition during in vivo bacterial infection. Cell Host Microbe. 2012;12(2):177–86.

10. Novick RP, Ram G. The Floating (Pathogenicity) Island: A Genomic Dessert. Trends Genet. 2016;32(2):114–26.

11. Box AM, McGuffie MJ, O’Hara BJ, Seed KD. Functional analysis of bacteriophage immunity through a Type I-E CRISPR-Cas system in *Vibrio cholerae* and its application in bacteriophage genome engineering. J Bacteriol. 2015;198(3):578–90.

12. O’Hara BJ, Barth ZK, McKitterick AC, Seed KD. A highly specific phage defense system is a conserved feature of the *Vibrio cholerae* mobilome. PLoS Genet. 2017;13(6):e1006838.

13. Seed KD, Lazinski DW, Calderwood SB, Camilli A. A bacteriophage encodes its own CRISPR/Cas adaptive response to evade host innate immunity. Nature. 2013;494(7438):489–91.

14. Wilkinson ME, Nakatani Y, Staals RHJ, Kieper SN, Opel-Reading HK, McKenzie RE, et al. Structural plasticity and *in vivo* activity of Cas1 from the type I-F CRISPR-Cas system. Biochem J. 2016;473(8):1063–72.

15. Richter C, Fineran PC. The subtype I-F CRISPR–Cas system influences pathogenicity island retention in Pectobacterium atrosepticum via crRNA generation and Csy complex formation. Biochem Soc Trans. 2013;41(6):1468–74.

16. Fagerlund RD, Wilkinson ME, Klykov O, Barendregt A, Pearce FG, Kieper SN, et al. Spacer capture and integration by a type I-F Cas1–Cas2-3 CRISPR adaptation complex. Proc Natl Acad Sci. 2017;114(26):E5122–8.

17. Rollins MF, Chowdhury S, Carter J, Golden SM, Wilkinson RA, Bondy-Denomy J, et al. Cas1 and the Csy complex are opposing regulators of Cas2/3 nuclease activity. Proc Natl Acad Sci. 2017;114(26):E5113–21.

18. Bellas CM, Anesio AM, Barker G. Analysis of virus genomes from glacial environments reveals novel virus groups with unusual host interactions. Front Microbiol. 2015;6(656).

19. Chénard C, Wirth JF, Suttle CA. Viruses Infecting a Freshwater Filamentous Cyanobacterium (Nostoc sp.) Encode a Functional CRISPR Array and a Proteobacterial DNA Polymerase B. MBio. 2016;7(3):e00667–16.

20. McKitterick AC, Seed KD. Anti-phage islands force their target phage to directly mediate island excision and spread. Nat Commun. 2018;9(2348).

21. Naser I Bin, Hoque MM, Nahid MA, Rocky MK, Faruque SM. Analysis of the CRISPR-Cas system in bacteriophages active on epidemic strains of *Vibrio cholerae* in Bangladesh. Sci Rep. 2017;7(1):14880.

22. Seed KD, Bodi KL, Kropinski AM, Ackermann H-W, Calderwood SB, Qadri F, et al. Evidence of a Dominant Lineage of *Vibrio cholerae*-Specific Lytic Bacteriophages Shed by Cholera Patiens over a 10-Year Period in Dhaka, Bangladesh. MBio. 2011;2(1):e00334–10.

23. Angermeyer A, Das MM, Singh DV, Seed KD. Analysis of 19 Highly Conserved Vibrio cholerae Bacteriophages Isolated from Environmental and Patient Sources Over a Twelve-Year Period. Viruses. 2018;10(6):299.

24. Koren S, Walenz BP, Berlin K, Miller JR, Bergman NH, Phillippy AM. Canu: scalable and accurate long-read assembly via adaptive *k*-mer weighting and repeat separation. Genome Res. 2017;27(5):722–36.

25. Walker BJ, Abeel T, Shea T, Priest M, Abouelliel A, Sakthikumar S, et al. Pilon: An Integrated Tool for Comprehensive Microbial Variant Detection and Genome Assembly Improvement. PLoS One. 2014;9(11):e112963.

26. Dalia AB, Lazinski DW, Camilli A. Identification of a Membrane-Bound Transcriptional Regulator That Links Chitin and Natural Competence in Vibrio cholerae. MBio. 2014;5(1):e01028–13.

27. Lazinski DW, Camilli A. Homopolymer tail-mediated ligation PCR: A streamlined and highly efficient method for DNA cloning and library construction. Biotechniques. 2013;54(1):25–34.

28. Langmead B, Trapnell C, Pop M, Salzberg SL. Ultrafast and memory-efficient alignment of short DNA sequences to the human genome. Genome Biol. 2009;10(3):R25.

29. Cock PJA, Antao T, Chang JT, Chapman BA, Cox CJ, Dalke A, et al. Biopython: freely available Python tools for computational molecular biology and bioinformatics. Bioinformatics. 2009;25(11):1422–3.

30. Barker A, Clark CA, Manning PA. Identification of VCR, a repeated sequence associated with a locus encoding a hemagglutinin in *Vibrio cholerae* O1. J Bacteriol. 1994;176(17):5450–8.

31. Staals RHJ, Jackson SA, Biswas A, Brouns SJJ, Brown CM, Fineran PC. Interference-driven spacer acquisition is dominant over naive and primed adaptation in a native CRISPR-Cas system. Nat Commun. 2016;7:12853.

32. Vorontsova D, Datsenko KA, Medvedeva S, Bondy-Denomy J, Savitskaya EE, Pougach K, et al. Foreign DNA acquisition by the I-F CRISPR-Cas system requires all components of the interference machinery. Nucleic Acids Res. 2015;43(22):10848–60.

33. Van Erp PBG, Jackson RN, Carter J, Golden SM, Bailey S, Wiedenheft B. Mechanism of CRISPR-RNA guided recognition of DNA targets in *Escherichia coli*. Nucleic Acids Res. 2015;43(17):8381–91.

34. Van Houte S, Ekroth AKE, Broniewski JM, Chabas H, Ashby B, Bondy-Denomy J, et al. The diversity-generating benefits of a prokaryotic adaptive immune system. Nature. 2016;532(7599):385–8.

35. Richter C, Dy RL, McKenzie RE, Watson BNJ, Taylor C, Chang JT, et al. Priming in the Type I-F CRISPR-Cas system triggers strand-independent spacer acquisition, bi-directionally from the primed protospacer. Nucleic Acids Res. 2014;42(13):8516–26.

36. Xue C, Seetharam AS, Musharova O, Severinov K, Brouns SJJ, Severin AJ, et al. CRISPR interference and priming varies with individual spacer sequences. Nucleic Acids Res. 2015;43(22):10831–47.

37. Mojica FJM, Díez-Villaseñor C, García-Martínez J, Almendros C. Short motif sequences determine the targets of the prokaryotic CRISPR defence system. Microbiology. 2009;155(3):733–40.

38. Redding S, Sternberg SH, Marshall M, Gibb B, Bhat P, Guegler CK, et al. Surveillance and Processing of Foreign DNA by the *Escherichia coli* CRISPR-Cas System. Cell. 2015;163(4):854–65.

39. Brown MW, Dillard KE, Xiao Y, Dolan AE, Hernandez ET, Dahlhauser S, et al. Assembly and translocation of a CRISPR-Cas primed acquisition complex. bioRxiv [Internet]. 2017; Available from: https://doi.org/10.1101/208058

40. Garneau JE, Dupuis M-E, Villion M, Romero DA, Barrangou R, Boyaval P, et al. The CRISPR/Cas bacterial immune system cleaves bacteriophage and plasmid DNA. Nature. 2010;468(7320):67–71.

41. Jinek M, Chylinski K, Fonfara I, Hauer M, Doudna JA, Charpentier E. A Programmable Dual-RNA–guided DNA endonuclease in adaptive bacterial immunity. Science. 2012;337(6096):816–21.

42. Makarova KS, Wolf YI, Alkhnbashi OS, Costa F, Shah SA, Saunders SJ, et al. An updated evolutionary classification of CRISPR-Cas systems. Nat Rev Microbiol. 2015;13(11):722–36.

43. Yosef I, Manor M, Kiro R, Qimron U. Temperate and lytic bacteriophages programmed to sensitize and kill antibiotic-resistant bacteria. Proc Natl Acad Sci. 2015;112(23):7267–72.

44. Bikard D, Euler CW, Jiang W, Nussenzweig PM, Goldberg GW, Duportet X, et al. Exploiting CRISPR-cas nucleases to produce sequence-specific antimicrobials. Nat Biotechnol. 2014;32(11):1146–50.

45. Crooks G, Hon G, Chandonia J, Brenner S. WebLogo: a sequence logo generator. Genome Res. 2004;14(6):1188–90.

46. Bawono P, Heringa J. PRALINE: A Versatile Multiple Sequence Alignment Toolkit. Methods Mol Bio. 2014;1079:245–62.

